# Key determinants of CRISPR/Cas9 induced inversions in tomato

**DOI:** 10.1101/2024.01.09.574821

**Authors:** Jillis Grubben, Gerard Bijsterbosch, Richard G.F. Visser, Henk J. Schouten

## Abstract

Inversions in chromosomes occur widely within plants and suppress meiotic recombination which can be beneficial or detrimental for plant breeders. Therefore, induction or reversion of inversions via CRISPR/Cas9 has been extensively researched recently. Extensive variation in inversion induction rates and sizes have been reported, from hundreds to several million base pairs. Here, we dissect the influential factors of inversion induction efficiency using CRISPR/Cas9. By using a fixed reference gRNA, we could directly correlate gRNA mutation frequency to inversion frequency and inversion size, of inversions up to 37.5 Mb in length in tomato. Our findings indicate that the least efficient gRNA is the bottleneck for inversion induction, with inversion size having no significant influence unless the inversions were larger than 1 Mb in size. For these huge inversions, the frequency dropped astoundingly, regardless of the gRNA cutting efficiencies. We hypothesize an *in planta* yet unknown variant of non-homologous-end-joining (NHEJ)-based repair which utilizes active transport of damaged chromosomal sections to dedicated repair sites in the cell nucleus, where repair is finalized. We propose that large inversions are formed less frequently because the transport of these segments to the repair sites may be hampered by their sheer size.

## Introduction

Genomic rearrangements, including inversions, are a significant source of genetic diversity among related species (Coughlan and Willis, 2019). An inversion is a type of genomic discontinuity in which a chromosomal segment is flipped to form the mirror image of the original segment (Hoffmann, *et al.,* 2004). Inversions can vary greatly in size, with reports of inversions ranging from tens of base pairs to hundreds of kilobase pairs (Wolters *et al*., 2015 and Zapata *et al*., 2016).

Inversion events can occur naturally and may become fixed in a population, potentially causing fitness effects, reduced meiotic recombination, and even speciation events (Ortiz- Barrientos et al., 2016). There are two proposed mechanisms for inversion formation: spontaneous excision and inversion of a genomic region between two transposons, followed by re-ligation into the genome (Gray, 2000); and simultaneous double-strand breaks in a chromosome followed by repair through canonical non-homologous end-joining (c-NHEJ) in the inverted orientation (Feschotte & Pritham, 2007). In meiosis, inversions may pair in heterozygotes through inversion loops, but a crossover in an inversion loop can result in the loss of large chromosomal sections, which is lethal to progeny (Huang & Rieseberg, 2020). As a result, crossovers in inversions are rare in organisms that are heterozygous for the inversion (Verlaan *et al*., 2011; Connallon & Olito, 2022). Consequently, it is very difficult for plant breeders to remove linkage drag from an inversion, if the gene of interest is located in that inversion.

Several resistance genes in wild tomato relatives are located within inversions when compared to the reference genome. Examples include the *Ty-2* gene on chromosome 11 in *Solanum habrochaites*, which confers resistance to tomato yellow leaf curl virus, and the *Mi- 1* gene on chromosome 6 in *S. peruvianum*, which provides resistance to root-knot nematodes (Hanson, Green, & Kuo, 2006; Milligan *et al*., 1998). These genes are located within inversions of 200 kb and 300 kb in size, respectively (Wolters *et al*., 2015; Seah, Telleen, & Williamson, 2007). These inversions prevent recombination with the homologous chromosomal segments in tomato varieties (*S. lycopersicum*) (Szinay, *et al.,* 2010). Consequently, all modern tomato varieties harbouring the *Mi-1* gene and/or the *Ty-*2 gene do carry linkage drag from the wild donors. This issue in plant breeding has not been solved yet, although it exists already for decades (Verlaan *et* al., 2011).

However, the linkage drag issue caused by inversions can be solved. Gene-editing technology has made it possible to induce two double-strand breaks (DSBs) at specific locations of a chromosome, using CRISPR/Cas (Ran *et al*., 2014). This can result in excision of the DNA fragment between the DSBs, which can be ligated back as the mirror image of the wild type via c-NHEJ, resulting in an inversion event.

The CRISPR/Cas9 system has been used to successfully induce targeted inversions in a variety of species (Zhang *et al.,* 2017; Xiao *et al.,* 2013; Schmidt *et al.,* 2019; Schwartz *et al.,* 2020; Lu *et al*., 2021). In a study by Zhang and colleagues (2017), the use of constructs expressing Cas9 together with two gRNAs resulted in the cutting out of DNA segments of up to 341 bp in size, with a frequency of 2.6% of these segments being reintroduced in the genome at the site of the DSBs in an inverted fashion. Schmidt *et al*. (2019) were able to make scarless heritable inversions in *Arabidopsis thaliana* of up to 18 kb in length using floral dip. Genome-editing methods that allow for the induction of genomic rearrangements such as inversions offer new opportunities for breeders. For example, the induction of inversions via CRISPR/Cas can re-shuffle promoter-gene combinations by cutting out DNA segments located between two promoter-gene borders on the same chromosome, as demonstrated in rice where a heritable inversion of 911 kb in size was induced (Lu *et al*., 2021). Schmidt *et al*. (2020) reverted a 1.1 Mb inversion in *A. thaliana*, allowing normal chromosome pairing during meiosis and enabling recombination to occur in the reverted region. This approach was also applied in maize, where an elite inbred line was used for the CRISPR/Cas-mediated reversion of a 75.5 Mb inversion (Schwartz *et al.,* 2020). Traditional plant breeding methods are not able to break this linkage drag.

Further, induction of inversions can prevent recombination between favourable genes that are physically bound together in a cluster on a chromosome, as demonstrated by Rönspies *et al*. (2022), who inverted 90% of chromosome 2 in *A. thaliana*, effectively suppressing crossovers in almost the entire chromosome. In summary, the use of genome- editing methods for inducing inversions offers a range of possibilities for plant breeders, including the re-shuffling of promoter-gene combinations, the ability to overcome linkage drag, and the fixing of favourable gene clusters.

Recent advances have enabled the generation of inversions of various sizes, ranging from kilobases to tens of megabases, in multiple plant species with no apparent size limit. However, the size range of inversions reported in these studies often falls within a single order of magnitude and presents a limited number of inversion events, making it difficult to determine the relative difficulty of generating large versus small inversions. Additionally, no studies have examined the relationship between inversion frequency and size. The goal of our research was to investigate the relationship between inversion length and frequency of CRISPR/Cas-induced inversions. For this purpose, we induced a series of inversions, ranging from 1 kb to 37.5 Mb in chromosome 6 of tomato. The latter inversion covered approximately three-quarters of the chromosome including the centromere. Our results showed that the gRNA cutting frequency was the most significant factor in determining induction frequency. Surprisingly, the inversion size had no significant impact on the frequency of induced inversions for inversions up to 1 Mb. However, inversions larger than 1 Mb in size had significantly lower frequencies, even at high gRNA-cutting activity. We discuss a model that may explain this result.

## Results

### Inversion induction in tomato protoplasts yields inversions of up to 37.5 Mb in size

We employed thirteen CRISPR/Cas9 constructs, each carrying two gRNAs, to generate inversions through protoplast pool transfection. One of the gRNAs induced a DSB at a consistent position on chromosome 6, while the other gRNA generated a cut at a predetermined distance from the fixed gRNA, also on the same chromosome (referred to as the ‘fixed’ and the ‘variable’ gRNA; Figure 1a, b; Supplementary Table 1). We aimed to generate inversions by cutting at both gRNA target sites within a nucleus, leading to the excision of the inter-DSB DNA region. The repair mechanisms would repair the DNA, potentially re-ligating the excised fragment ends at the opposite DSB site to create an inversion. We incorporated the green fluorescent protein (GFP) gene into our constructs to estimate transfection efficiency. After tomato protoplast transfection, inversion-specific primer pairs were employed to amplify regions at both inversion ends, verifying induced inversion presence (Figure 1a).

**Figure 1.**
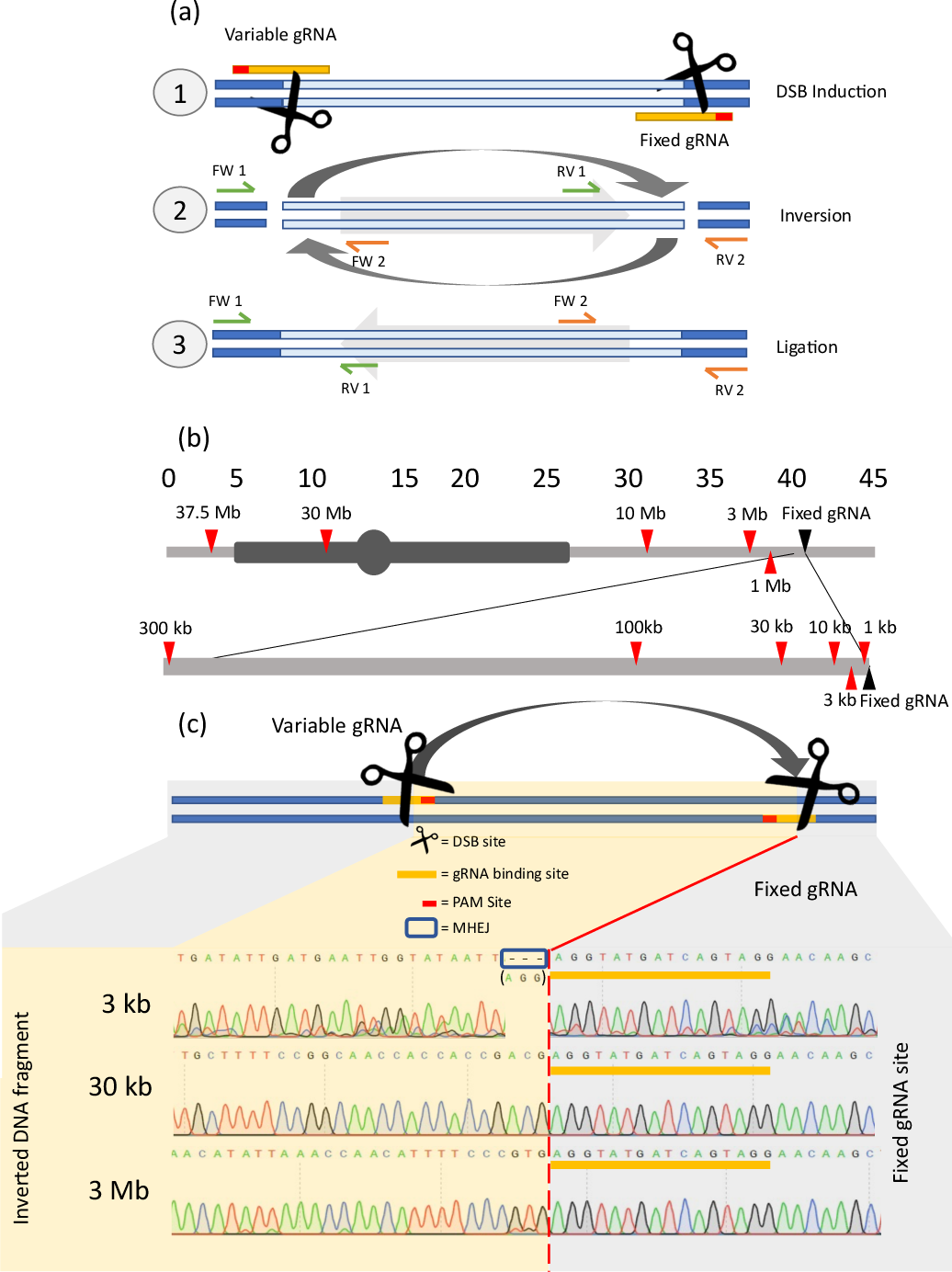
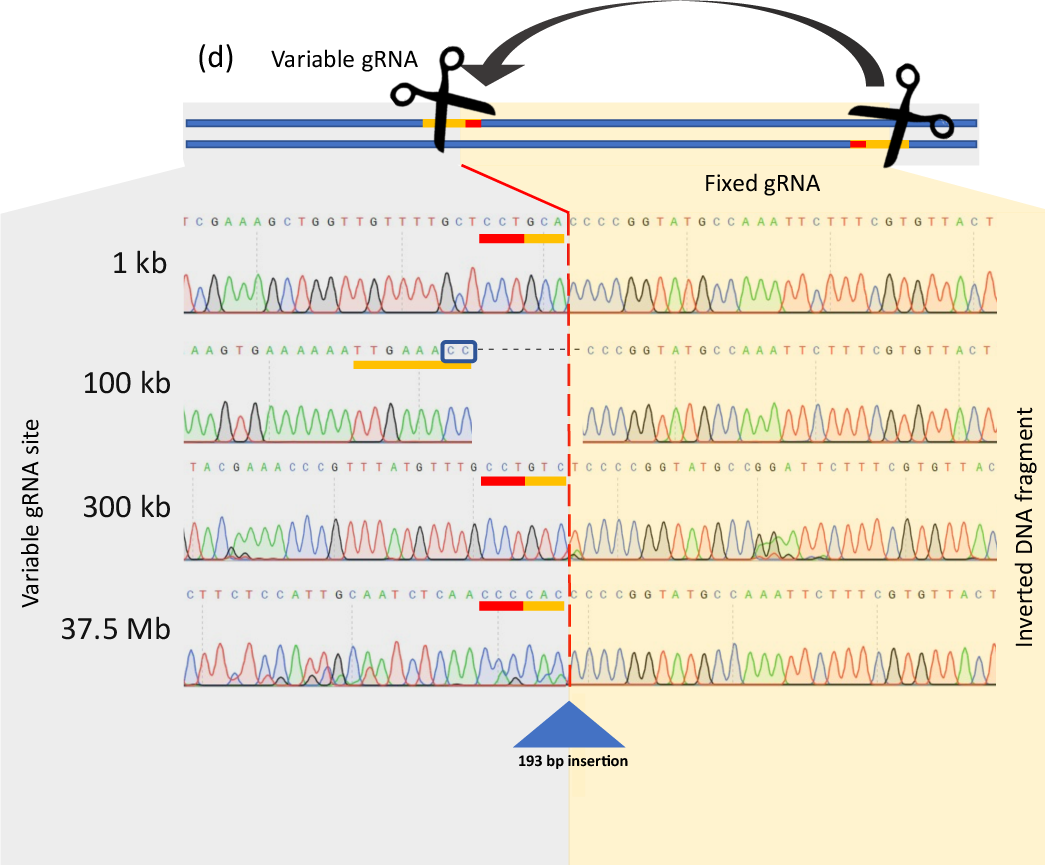
Cartoon showing (a) the gRNAs used to induce DSBs for inversion induction, and the primer pairs utilized to generate amplicons in the case of inversion events; (b) chromosome 6 including the cutting sites of the fixed gRNA (black triangle), and of the variable gRNAs (red triangles). The distances of the DSBs from the variable gRNAs to the DSB from the fixed gRNA are shown. The heterochromatin and the centromere are displayed in dark grey; (c) Sanger sequences of a subset of induced inversions detected at the fixed DSB site. Ligated inversions of 3 kb, 30 kb, and 3000 kb are shown as examples. (d) Sanger sequences of inversions detected at the distal gRNA cutting site. Grey and yellow surfaces indicate the WT DNA region, and the inverted DNA fragment, respectively. The red spaced line indicates the expected gRNA cutting site. Blue triangles indicate insertions, blue boxes indicate putative micro-homology-mediated NHEJ events, yellow bars indicate gRNA binding sites and red bars indicate PAM sites.

Sanger sequencing of inversion ends showed that we successfully generated inversions ranging from 1 kb to 37.5 Mb in size (Figure 1c, d; Supplementary Figure 1). We examined the predicted DSB sites at the ends of the inversions and found multiple inversion events directly at the predicted cutting sites, indicating perfect repair. In most remaining cases with imperfect repair, small fragments of three, six, or nine base pairs were deleted at the DSB site. For inversions of 3 kb in size, we detected no seamless re-ligation at the ‘fixed’ gRNA target location. Instead, we observed either a three-base-pair deletion or a six-base-pair deletion. These small deletions might have been the result of microhomology-mediated end-joining (MMEJ), a process in which small sections of DNA are removed during DNA repair.

### Addressing Transfection Efficiency Variability

Next, we wanted to determine the impact of inversion size on the frequency of inversion generation. To analyse the relationship between inversion size and frequency, we transfected CRISPR/Cas9 constructs into protoplast pools and employed crystal digital PCR (cdPCR) to identify inversion events. In parallel, on these samples we utilized Hi-Seq sequencing to detect mutations at the double-strand break (DSB) sites at both the ’fixed’ and ’variable’ gRNA target sites. This dual approach allowed us to correlate the frequency of CRISPR/Cas9-induced inversions with the mutation induction frequency, as illustrated in Figure 2a.

**Figure 2.**
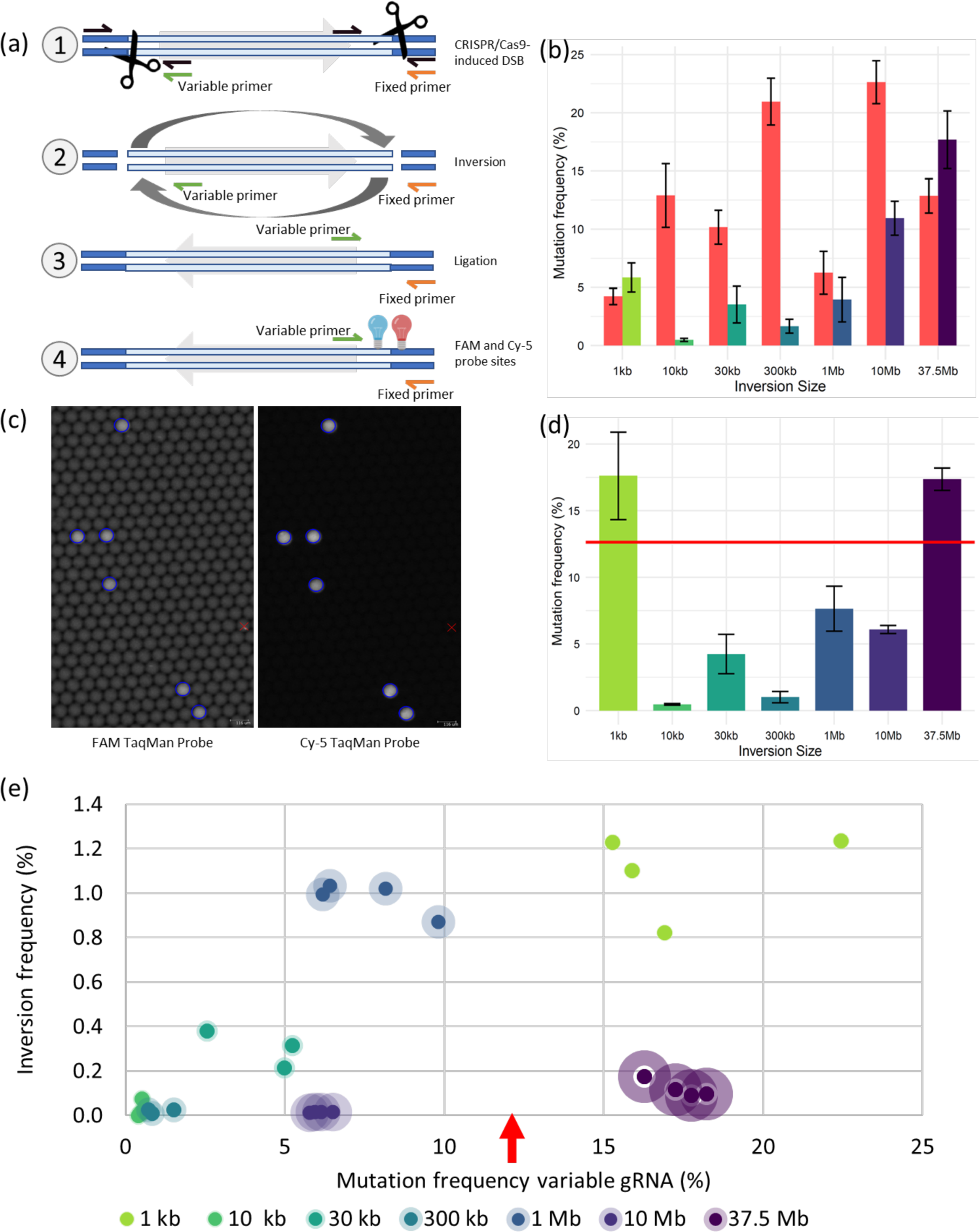
**a**, Schematic illustration of the orientation of fixed and variable cdPCR primers, and the binding sites of two TaqMan probes. The dark blue bars represent wild-type genetic regions and the light blue bars represent the DNA region to be inverted. The grey arrows show the relative orientation of the to-be-inverted region, and the variable and fixed primers are shown in green and orange, respectively. Black arrows indicate Hi-seq sequencing primer pairs for non-inversions. The figure shows the different phases in the experiment: (1) Before DSB induction, with the cutting positions of the variable gRNA (left) and the fixed gRNA (right) represented by scissors; (2) After DSB induction; (3) After inversion induction; (4) The binding position of the FAM TaqMan probe on the left inversion junction and the Cy-5 probe binding on the right side of the inversion junction are depicted with blue and red light bulbs, respectively. **b,** Comparison of mutation frequencies for variable and fixed gRNAs across various inversion sizes. The variable gRNA mutation frequencies are represented by a gradient colour scale (green to purple), while the fixed gRNA mutation frequency is shown in red. **c**, cdPCR droplet crystal matrix structure showing 1 Mb inversion events in cdPCR reaction volumes. Droplets encircled in blue indicate inversion events in the FAM (left) and Cy-5 (right) detection channels. Red crosses indicate droplets excluded from analysis by the cdPCR software. **d**, Variable gRNA mutation frequencies (%) after correcting for the transfection efficiency using the average mutation frequency of the fixed gRNA. The red vertical line represents the average mutation frequency of the fixed gRNA. **e**, Relationship between the inversion frequency at the vertical axis versus the mutation frequency of the variable gRNA at the horizontal axis. Both parameters are corrected for transfection efficiency, using the mutation frequency at the fixed gRNA site as a reference. Colour-labelled data points indicate biological replicates of samples transfected with inversion-inducing CRISPR/Cas9 constructs. The transparent halo around the data points indicates the size of the induced inversion. The red arrow indicates the average fixed gRNA mutation frequency of all samples.

We noted that variability in CRISPR/Cas9 plasmid delivery efficiency could affect inversion rates. Although GFP-induced fluorescence could correct for plasmid delivery variability, it is an indirect and potentially inaccurate measure. Our solution was a novel normalization strategy using an internal standard gRNA, the ‘’fixed’’ gRNA, which accounted for variations in transfection efficiency across all samples. By analysing the mutation induction of the internal standard gRNA target site, we could directly compare the efficiency of CRISPR/Cas9 between samples. This strategy enabled us to correct both the inversion frequency and the variable gRNA mutation frequency as shown in Figure 2b, d with the red horizontal line. Analysis showed the mutation ratio of ’variable’ to ’fixed’ gRNA remained stable across replicates (Supplementary Figure 3), indicating consistent mutation frequencies when co-expressed, thus validating their use as internal controls. Consequently, the factors that could potentially influence inversion frequency are the cutting efficiency of the ‘variable’ gRNA, the size of the inversions, and other unknown factors.

### Influence of gRNA Efficiency and Inversion Size on Induced Inversion Frequencies

Our results indicate that 1 kb inversions occurred in 1.1% of genomes analysed by cdPCR (Figure 1a), while larger inversions of 1 Mb occurred at a rate of 0.98%. Inversions of 30 kb were detected at a rate of 0.3%, and the largest inversion we attempted to generate, spanning 37.5 Mb, had a prevalence of 0.1% in our cdPCR set-up. Inversion sizes of 10 kb, 300 kb, and 10 Mb were rare, occurring in less than 0.05% of the genomes screened by cdPCR. Although some of the inversion sizes we detected were less frequent, they cannot be disregarded as random background noise. Our control experiments, in which protoplast pools were transfected with constructs lacking CRISPR/Cas9, yielded no inversion events in over 500,000 droplet-partitioned cdPCR reactions, proving the specificity and sensitivity of our detection method in identifying low inversion copy numbers per sample.

The broad range of inversion frequencies that we were able to induce was correlated with the frequency at which the chromosomes were cut. Although the ‘fixed’ gRNA was efficient in inducing mutations, the ‘variable’ gRNAs showed a range of mutation induction efficiencies, as 1 kb and 37.5 Mb variable gRNAs outperformed the ‘fixed’ gRNA, whereas the 3 kb, 100 kb, 3 Mb, and 30 Mb samples did not show any induced mutations at the CRISPR/Cas9 target sites, indicating that these gRNAs were not effective in inducing DSBs. Consistently, for the latter four groups no inversions were detected via cdPCR (Figure 2 b; Supplementary Figure 2).

When we plotted inversion induction and gRNA mutation frequencies against each other, we observed a positive trend (Figure 2e). Higher gRNA mutation induction frequencies were associated with more inversion events, and this trend was followed for inversions up to 1 Mb in size. Apparently, in this range, the frequency of DSB induction of the ‘variable’ gRNA was the bottleneck.

For inversions up to 1 Mb in size, we found that the mutation frequency of the variable gRNA was the main driver of inversion induction, irrespective of the mutation size (Figure 2 e). This linear trend could be followed up to a mutation frequency of 10 %, after which we saw a plateauing effect. Above this mutation frequency, the number of induced inversions did not increase anymore, as likely the ‘fixed’ gRNA cutting frequency became the limiting factor (Figure 2 d and e, red line and red arrow, respectively). For the 1 kb and 37.5 Mb samples, the ‘variable’ gRNA performed better than the ‘fixed’ gRNA in terms of mutation induction (Figure 2e, red arrow). Therefore, the ‘fixed’ gRNA was the bottleneck for inversion induction in these samples.

Strikingly, the two largest inversions we generated did not follow this trend. Instead, inversions of 10 Mb and 37.5 Mb had a lower prevalence of inversion events, even when having a relatively high ‘variable’ gRNA mutation induction frequency. It appears that the size of the inversion has now become a limiting factor. However, when analysing inversion sizes ranging from 1 kb to 1 Mb, we did not observe a significant impact of inversion size on inversion frequency (Figure 2e). It was only when the inversions were extremely large, specifically 10 Mb or 37.5 Mb, that the size of the inversion appeared to play a critical role.

### Validation of Complete Inversions and Comparison of Detection Methods

We performed new transfections in protoplasts to confirm the occurrence of complete inversions at both DSB ends in a single DNA molecule. Using PacBio Sequel II sequencing, we analysed the complete inversion region of 1 kb and 3 kb samples, confirming the presence of complete inversions for both sizes (Figure 3 a). The inversion frequency for 1 kb inversions was found to be 0.41%, while inversions of 3 kb were rare, with only one detected inversion event detected via PacBio sequencing. The low inversion induction frequency of the 3 kb inversion was likely due to the inefficiency of the gRNA at the 3 kb site (Figure 3 b, green bars). Comparing PacBio and cdPCR inversion detection methods, we found their results highly consistent (Figure 3 b), suggesting that analysing one junction via cdPCR and Hi-Seq sequencing can yield comparable results to full inversion sequence analysis.

**Figure 3.**
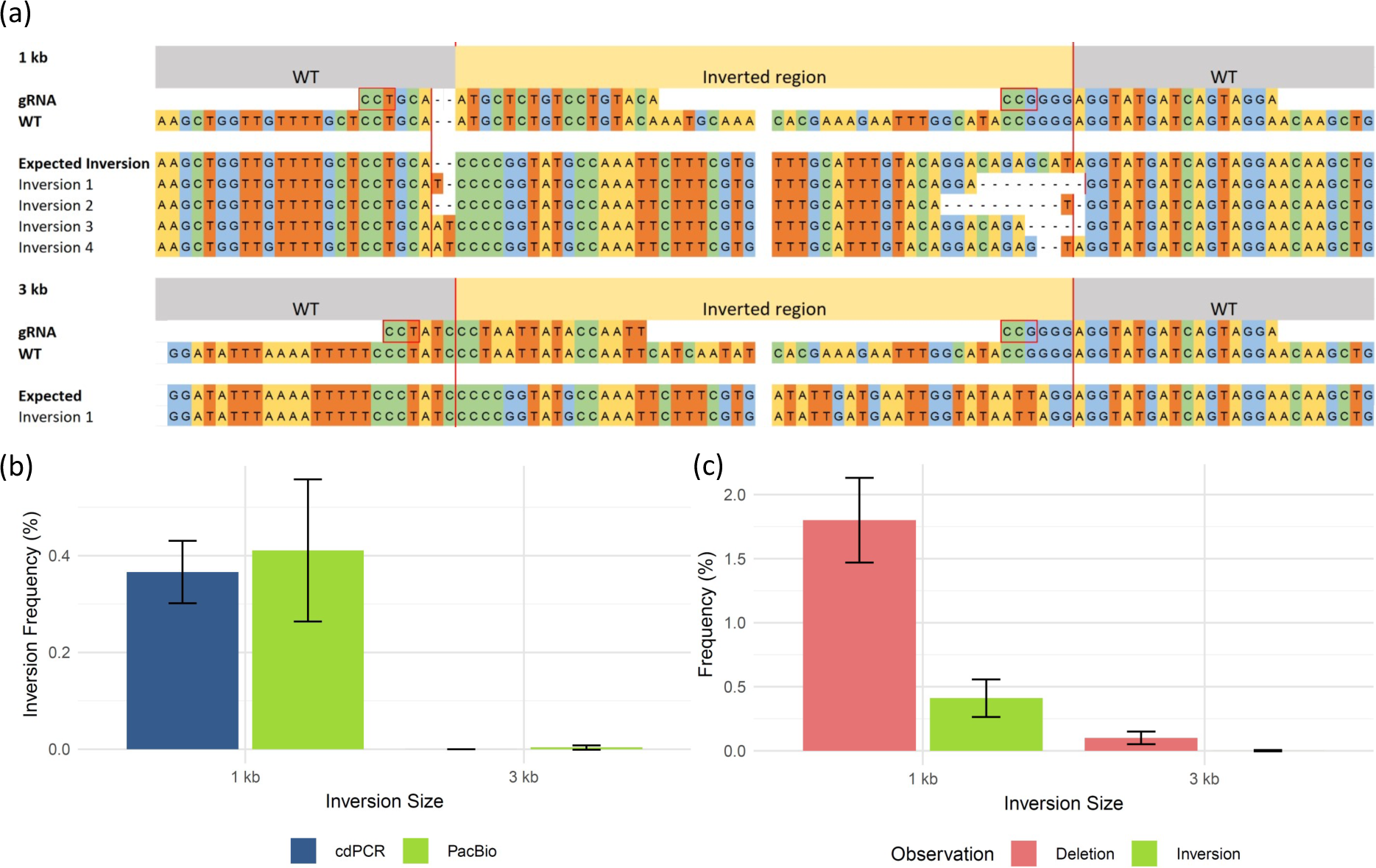
a) PacBio Sequel II amplicons displaying a selection of whole inversion sequences for 1 kb and 3 kb inversions. The red lines represent the expected cutting sites of the variable (left) and the fixed (right) cutting sites. Per inversion length, the gRNA sequence, the WT sequence, the expected inversion sequence, and the amplicons proving the inversion events are shown. Red boxes indicate PAM sites. Grey and yellow surfaces represent the WT DNA segments and the inverted DNA region, respectively. b) Entire inversion events of 1 kb and 3 kb in size detected via PacBio Sequel II sequencing (green) and via cdPCR + Hi-Seq (blue). The Y-axis shows the inversion frequency. c) Frequency of detected large deletions (red) and inversions (green) in the PacBio Sequel II Sequencing reads for 1 kb and 3 kb samples. Error bars represent standard deviations.

### Quantification of large intra-double strand break deletions and Inverted ligation with the homologous chromosome

Our analysis of PacBio Sequel II sequencing data revealed the occurrence of large deletions resulting from the removal of the region between the two induced DSBs on the same chromosomal molecule, with either the 1 kb or 3 kb inversion-inducing constructs (Figure 3 c). These pairs of DSBs can potentially lead to three outcomes: re-ligation of the broken DNA strands with or without small mutations at the DSB sites, generation of an inversion, or ligation of DSB sites, excising the intervening region. We explored a potential correlation between these large deletion events and inversion induction frequency. After adjusting for transfection efficiency using the ’fixed’ gRNA mutation induction rate, we quantified large deletion frequency. In the samples transfected with the 1 kb construct, we found that intra-DSB deletions occurred approximately four times more frequently than inversions. The inversion- to-deletion ratio in 3 kb samples could not be determined due to insufficient inversion events.

## Discussion

### Why did the frequency of inversions not depend on inversion size, unless the inversions were very large?

The most striking finding of our work is that the frequency of induced inversions did not depend on the size of the inversion. Specifically, unless exceeding 1 Mb, the size of the inversion seemed to have minimal impact. Instead, the primary factor for the frequency of induced inversions was the cutting frequency at the two gRNA sites. The latter relationship is evident, as induction of inversions can occur only when the two DSBs are induced simultaneously at the two target sites. For inversions to occur, the incorrect cutting ends must be brought close together for ligation. We expected that the inversion frequency would decrease at increasing inversion size, due to the increasing physical distance. However, excluding the very large inversions, this was not observed. Our findings are in line with experiments performed in yeast where HO endonuclease was used to generate inversion induction frequencies of 13 kb to 358 kb in size (Sunder & Wilson, 2019). Sunder and Wilson (2019) observed no significant difference in inversion occurrence between the induced inversions and concluded that inversion size did not negatively impact inversion induction frequency. While our results are coherent with the abovementioned findings, our conclusion is not. Namely, we generated inversions larger than 358 kb and for our inversions that were larger than 1 Mb, we observed a striking decrease in inversion induction, even when gRNA cutting efficiency was high (Figure 2e).

Inversions arise from NHEJ-mediated repair. For HR-based repair in multiple higher eukaryotes active transport to repair compartments in the nucleus has been described (Caridi et al., 2018; Hirakawa & Matsunaga, 2019; Meschichi *et al.,* 2022; Meschichi & Rosa, 2023; Schrank *et al.,* 2018). Still, understanding of DSB relocalization in NHEJ-mediated repair remains elusive. Following DSB formation, Ku70/80 complexes that initiate NHEJ-mediated repair rapidly associate around the DSBs and recruit other factors that stabilize the DNA (Zahid *et al.,* 2021; Zhao *et al.,* 2020). This sequence of events attempts to rapidly begin the NHEJ repair process while keeping the DSBs nearby, which would hint at *in situ* repair rather than at active transport to repair compartments. However, the study by Sunder and Wilson (2019) reported that whether the DSBs were initially close together or far apart in the genome before cleavage did not change how often they were repaired in a way that would cause inversions within the chromosome. This would hint at a mechanism at play that brings the DSBs in close proximity to each other after damage. Lobrich and Jeggo, (2017) distinguish fast and delayed forms of NHEJ repair, where the first is characterized by an attempt of the cell to rapidly stabilize and repair the DSB via the established c-NHEJ mechanism. When rapid c-NHEJ fails, in cases of complex or multiple DSBs, a slower c-NHEJ variant might be employed, involving more complex DSB processing. Wilson and Sunder (2020) further hypothesize the possibility of active transport of DSBs to dedicated repair sites, particularly in the context of this slower repair mechanism. The pervasive DSB induction by CRISPR/Cas9 at the target sites could initiate persistent repair stimuli to the cell, which might be interpreted as failure to repair the target site. This is backed by findings in yeast where persistent DNA damage leads to the migration of damaged DNA to the nuclear periphery for repair (Oza *et al.,* 2009). Such relocation could explain the absence of a relationship between the size of an inversion and its induction rate. After relocation, the DSBs are brought into proximity and are repaired, making the original spatial distance between the DSBs irrelevant.

We propose that plants may possess dedicated DSB repair centres where NHEJ-mediated repair is facilitated for sites that face persistent damage. Broken DNA segments would be transported to these sites, where inversions occur when the wrong ends are illegitimately ligated together (Figure 4). For larger DNA segments, we propose that physical constraints such as entanglement may hinder their tethering from two DSB sites to repair sites. This could significantly impede the transportation and subsequent repair process, leading to a reduced frequency of repair, and consequently, a lower incidence of chromosomal inversions (Figure 4). Unrepaired DSBs could lead to loss of chromosomal sections followed possibly by cell death.

**Figure 4.**
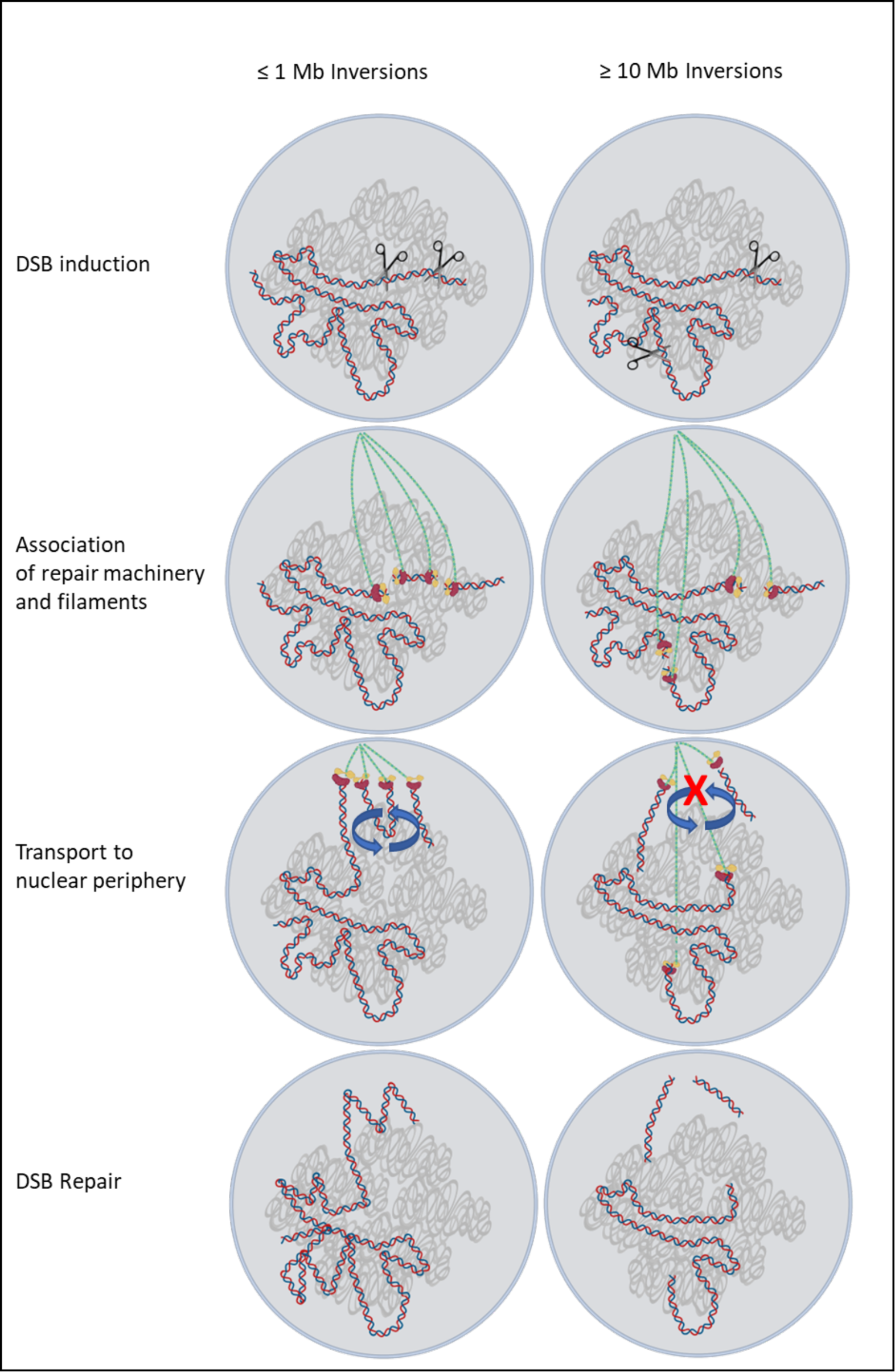
Comparative illustration of a model for the post-DSB induction process for inversions smaller and larger than 1 Mb in case of active transport of DSBs to the nuclear periphery. From top to bottom, presented are the CRISPR/Cas9 target sites, followed by the recruitment of the DSB repair machinery of the cell to these sites, which includes the generation of long filaments (depicted as green strings) extending from the DSB sites to the nuclear periphery. The subsequent step involves the active translocation of DSBs along these filaments to the nuclear periphery. The potential for inversion is indicated by blue arrows, with the crossed-out red lines suggesting a lower likelihood of such events for inversions larger than 1 Mb. For DNA segments larger than 1 Mb, we propose that physical limitations, such as entanglement, might impede their ability to move from two sites of DNA breakage to sites for repair, reducing the frequencies of very large inversions.

Sunder and Wilson (2019) observed that small and large inversions occurred with similar frequencies. They did not observe the frequency drop of their largest inversions which were 358 kb in size. This agrees with our findings, as we also did not find a decrease in inversion induction frequency up to 1 Mb sized inversions. It was only above the 1 Mb threshold, when we observed the strong decrease in inversion induction frequency. Sunder and Wilson did not induce Mb-sized inversions. The yeast genome is approximately 100 times smaller than that of the tomato (Goffeau *et al.,* 1996; Gao *et al.,* 2019). In the more extensive tomato genome, transported DNA segments could encounter greater structural complexity, potentially increasing the likelihood of entanglement during their transport to repair sites. Notably, except for one, all our induced inversions were paracentric, indicating no significant centromeric effects on inversion induction.

### The cutting frequency of the least efficient gRNA site was the most important factor for the inversion frequency

Reported efficiencies and sizes for CRISPR/Cas-induced inversions vary across different plant species. Zhang *et al*. (2017) achieved a 2.6% efficiency for a 341 bp inversion in *A. thaliana*, and Schmidt *et al*. (2019) noted about 1% for 2.9–18 kb inversions. In rice, a 911 kb inversion occurred in three out of 259 calli, and in maize, a 75.5 Mb inversion was found in two out of 1,500 embryos (Lu *et al*. 2021; Schwartz *et al*. 2020). Intrigued by the diverse sizes of inducible inversions, we investigated how CRISPR/Cas9 cutting frequency affects inversion sizes in plant genomes.

As shown in Figure 2e, induced inversions increased proportionally with the mutation frequency of the variable gRNA, up to 10 %. However, beyond this threshold, the inversion frequency reached a plateau. This plateau may be caused by the limiting cutting frequency of the fixed gRNA, whose mutation frequency was 12.6 % (as indicated by the orange arrow in Figure 2e). We observed that constructs where variable gRNAs were unable to induce mutations also failed to generate inversions. We, therefore, conclude that the frequency of induced inversions is primarily determined by the cutting efficiency of the least efficient gRNA. This conclusion aligns with the results of Schmidt et al. (2019), who found that samples with higher double-strand break (DSB) frequencies generated more inversion events.

### A standard gRNA for correction of efficiency of plasmid delivery into protoplasts

We used a ’fixed’ gRNA as an internal control to correct for plasmid delivery efficiency into protoplasts. This method could be preferable to traditional gRNA efficiency estimation techniques as it offers a consistent normalization baseline that allows for direct measurement of the cutting efficiency against a known standard. Moreover, this method removes the need for indirect measures of delivery efficiency, such as GFP-positive cell counting via microscopy. Another field of application of using a reference gRNA is comparison of gRNAs. It is common practice to test gRNAs in protoplasts to verify their efficiency, before utilizing them in more laborious stable transformation experiments. By introducing an internal standard gRNA, it is possible to directly compare the mutation induction efficiency of a tested gRNA to a known standard, greatly improving good gRNA selection. This opens up the possibility also to study the effects of sequence or genomic context in relation to gRNA efficiency, as the internal standard gRNA can be utilized for normalisation.

Our data show that the mutation frequency ratio between a variable gRNA and the internal standard gRNA remains stable between samples treated with the same construct, underscoring the reliability of using a gRNA as a control (Figure 2 d; Supplementary Figure 3). Nonetheless, it is important to note that potential drawbacks, such as translocations and competition among gRNAs for binding to the Cas protein, may also affect the results. We recommend utilising the internal standard gRNA that targets a genomic site on a separate chromosome from that of the target gene, to avoid the formation of intrachromosomal deletions or inversions, and to use constructs that contain both the target gRNA as well as the internal standard gRNA in one.

### Two methods for quantifying inversions

Using cdPCR we screened for inversions at one inversion border. It is possible that the detected inversions were induced at the site we screened for, but not at the other site. Schmidt *et al*. (2019) also used a digital PCR method to screen for inversions but used probes at both inversion borders, finding no significant differences in inversion events at the two DSB sites. To verify that complete inversion events were captured in our study, we amplified the region that contained the inversion using PacBio Sequel II sequencing. Our two distinct approaches for detecting inversions, applied to different protoplast pools, resulted in highly similar results after normalization with our internal standard gRNA method. This uniformity across methods showed their reliability, with the added benefit of PacBio Sequel II sequencing that allowed for the confirmation of induction of complete inversion events. PacBio sequencing had a size limitation for inversions larger than 3 kb, whereas the cdPCR method did not have this limitation. Overall, our findings demonstrate that both cdPCR and PacBio sequencing are effective methods for detecting inversion events, but cdPCR can be used too for large inversions that are outside the range of PacBio amplicon sequencing.

### Large deletions

When a chromosome undergoes two simultaneous DSBs, it faces three potential repair paths: re-ligation of the excised fragment with potential minor indels; excision of the intervening section resulting in a major deletion; or an inversion event. This study concentrates on the latter scenario. However, PacBio Sequel II sequencing also revealed large inter-DSB deletions, indicating the second option. We performed PacBio sequencing of 1 kb and 3 kb fragments and found these large deletions in all samples of both sizes. Deletions of the 1 kb size were four times more frequent than inversions, consistent with prior observations. Schmidt *et al*. (2019) reported ratios ranging from 1.4 to 3.5, whereas Zhang *et al*. (2017) observed a ratio of 10, and Liu *et al*. (2023) identified deletion-to-inversion ratios varying between 1.3 and 6.6 in *A. thaliana* and between 1.3 and 14.3 in rice. Our results were obtained in protoplasts that were not undergoing cell division. Loss of large chromosomal fragments after cell division may be lethal for daughter plant cells.

## Conclusion

This study enriches our understanding of chromosomal inversion processes in plant cells induced by CRISPR/Cas. We found that the induction efficiency of chromosomal inversions is largely dependent on the activity of the least effective gRNA rather than on the size of the inversion, with the latter becoming a significant factor only when inversions exceed 1 Mb. This observation points to a distinctive size-dependent variation in inversion dynamics, that might be caused by the directed transport of DSB sites to dedicated repair foci, a phenomenon previously described for HDR-based repair, but not for NHEJ. By utilizing a novel method involving a standard gRNA that was responsible for one of the two DSBs that lead to inversions, we were able to dissect the dynamics of CRISPR/Cas9-dependent inversion induction. The use of an omnipresent standard gRNA allowed us to normalize plasmid delivery by correcting for the mutation frequency at the standard gRNA site, instead of indirect proxies such as GFP markers.

## Materials and Methods

### CRISPR/Cas9 constructs

For the induction of targeted inversions we created for each inversion length a CRISPR/Cas9 construct which contained two gRNAs targeting positions on the ch06: A “Fixed gRNA” with an earlier observed high mutation induction frequency was used in all constructs, while the other gRNA was variable, allowing for the induction of inversions of different lengths (Supplementary table 1). For the variable gRNA target regions with a distance to the fixed gRNA of approximately 1 kb, 3 kb, 10 kb, 30 kb, 100 kb, 300 kb, 1 Mb, 3Mb, 10 Mb, 30 Mb, and 37.5 Mb were selected, using CRISPOR (http://crispor.tefor.net/). gRNAs were synthesized (MacroGen Europe) and *BsaI* restriction sites were added via PCR. The gRNAs were inserted into LVL1 Golden Gate vectors containing the *U6-26* promoter via cut-ligation using *BsaI* and T4 ligase and subsequently these gRNAs were recombined with LVL 1 constructs containing pUbi::Cas9, NosP:: NPTII, and pCsVMV::GFP. Construct components were shown in Supplementary table 2. Plasmids were isolated using the Plasmid Midi Kit (Qiagen).

### Plant material and growth conditions

*S. lycopersicum* cv. Moneymaker seeds were sterilized in 1% NaClO for 20 minutes and washed in sterilized MilliQ. Seeds were sown on germination medium (½ MS including vitamins (Duchefa), 3% sucrose and 0.8% Daishin agar, pH 5.8) in sterile tissue culture vessels (OS140BOX/green filter, Duchefa), containing 10 seeds per vessel. Plants were grown *in vitro* at 24°C with a relative air humidity of 60% and a light intensity of 150 Wm^2^ in propagation containers for one month.

### Protoplast generation

Protoplast transfection was performed using a protocol adapted from (Maas & Werr, 1989). Approximately 20 small leaves from one month old seedlings were harvested and placed abaxial side down in a petri dish containing 10 mL digestion solution (0.4M mannitol, 20 mM MES, 20 mM KCl, 10mM CaCl2.2H2O, pH 5.7). Leaves were cut in a fine feather-like pattern, from the midrib to the edge of the leaves. The digestion solution was removed and subsequently 20 mL freshly prepared digestion solution containing 1% cellulase R10 and 0.3% Macerozym R10 enzymes (Duchefa) was added onto the leaves using a syringe with a 20 μm filter. Leaf material was incubated for digestion of the cell wall for 17 hours in the dark at 25°C. The plant material was gently swirled, after which the protoplast suspension was transferred through a Falcon 100 μm cell strainer (Corning) and carefully collected into a 50 mL tube using a 25 mL serological pipette. 10 mL of W5 washing buffer (154mM NaCl, 125mM CaCl2.2H2O, 5mM KCL and 2mM MES, pH5.7) was added to the petri dish and the dish was shaken 30 times to release additional protoplasts. The 50 mL tube containing the cell suspension was centrifuged at 100 RCF for 3 minutes at room temperature to pellet the protoplasts. Break and acceleration of the centrifuge were set at 5, Eppendorf 5810R, after which the supernatant was discarded. The protoplasts were resuspended in 10 mL fresh W5 and centrifugated again. Protoplasts were resuspended in 10 mL MMg solution (0.4M Mannitol, 15mM MgCl2 and 4mM MES, pH 5.7), centrifuged for 3 minutes at 100 RCF, and resuspended in 10 mL MMg. The protoplast density was estimated using a haemocytometer (Fuchs-Rosenthal) and diluted with MMg solution to a density of 1 million protoplasts per mL.

### Transfection

Per transfection, 10 μg of plasmid containing a CRISPR/Cas construct (final volume adjusted to 20 μL MilliQ) was added to a 2 mL Eppendorf tube. Using a 1 mL pipet tip with 0.5 cm of the bottom cut off, 200 μL protoplast suspension was pipetted on the plasmid, followed by the addition of 200 μL freshly prepared PEG solution (4.0 g PEG-4000 (Fluka), 3.0 mL MilliQ, mL 0.8M mannitol solution and 1.0 mL 1M CaCl2 solution). The suspension was gently but thoroughly mixed and incubated for 15 minutes after which 500 μl of W1 buffer (0.5M Mannitol, 20mM KCL and 4mM MES, pH5.7) was added droplet-wise to the suspension before continuing to the next sample. Another 500 μL of W1 was added to each sample following centrifugation at 200 RCF for 3 minutes. The supernatant was removed by pipetting and 1 mL of W1 was added to the protoplasts as described above. Samples were centrifuged, supernatant was removed, and protoplasts were resuspended in 150 μL W1. Protoplasts were incubated in the dark for 48 hours at 25°C. Protoplast transfection efficiency was estimated by estimating the percentage of fluorescing protoplasts using fluorescence microscopy. Supernatant was removed after centrifugation at 200 RCF for 60 seconds, and protoplast material was stored at -20°C.

### Protoplast DNA isolation for cdPCR and Hi-Seq sequencing

Protoplast DNA was isolated using the NucleoMag 96 Plant DNA-isolation kit (BioKé), following the provided manual DNA isolation protocol. DNA was eluded in 50 μL MilliQ. Impurities from the DNA-isolation kit that can interfere with the crystal matrix generation in the cdPCR system, were removed by NaCl/ethanol-based washing of the DNA samples. Per DNA sample 5 μg molecular biology grade glycogen (ThermoFisher), 5 μL 5M NaCl and 55 μL isopropanol were added and stored at -20°C for 24h after which the samples were thawed and centrifuged at 20.000g (4°C) for 1 hour. Supernatant was removed, followed by addition of 100 μL 70% ethanol and centrifugation at 20.000g (4°C) for 30 minutes. The supernatant was discarded, and DNA was dissolved in 15 μL Milli-Q. DNA concentrations were measured using the Qubit High Sensitivity kit (Invitrogen).

### Inversion detection using Sanger sequencing

Isolated DNA was enriched for inversion events at both DSB sites in separate PCR reactions using Phusion polymerase (ThermoFisher). Per inversion DSB border, two primers were utilized which both had a reverse orientation in wild-type DNA and therefore yielded no amplicons, while in DNA templates containing an inversion event, one of the primers served as forward primer, thus allowing for amplification of inversion events at the DSB location. For revealing the sequences of individual inversion border sequences, the amplicons were cloned into *Escherichia coli* using Golden Gate cloning. The plasmids containing the inversion borders were isolated from cell cultures originating from single colonies, using the QIAprep Spin Miniprep Kit (Qiagen). Inversion borders were analysed using Sanger sequencing (Macrogen Europe).

### Quantification of inversions using cdPCR. Primer and probe design

The cdPCR system from Stilla (Naica Geode and Naica Prism3) was used for the quantification of inversions. This system is based on droplets with uniform sizes called crystals, in which inversion-specific amplification can occur, leading to fluorescent signals from the droplets that harboured the inversion. For this purpose, inversion-specific primers were designed (Supplementary table 3). All primer pairs share the same ‘fixed primer’, while the other primer varied, depending on the used ‘variable gRNA’. These ‘fixed’ and ‘variable’ primers both have a reverse-orientation affinity to wild-type DNA, and thus do not yield amplicons in a PCR reaction. However, in the case of an inversion, the DNA segment including the primer binding site has flipped, allowing this primer to serve as a forward primer which will be in close proximity to the reverse “fixed” primer, enabling PCR-based amplification of the inversion boundary site (Figure 4).

TaqMan probes were used to detect inversion events in the individual crystals in the cdPCR system (Supplementary Figure 3). A FAM-labelled probe was designed to bind at one side of the inversion border, whereas a Cy-5 probe targeted the other side of the inversion boundary (Figure 2a). In case of inversions, both Taqman probes should emit their signals. For ensuring the primer pairs and probes worked properly in the cdPCR system, we ordered a synthetic stretch of DNA sequences of the expected inversion (g-blocks; Integrated DNA Technologies) and verified the reliability of the primer-probe combinations, and performed spill over tests between the FAM and the Cy-5 detection channels.

### cdPCR protocol

The cdPCR reaction mixtures were prepared using 5 μL of Perfecta Multiplex qScript ToughMix 5X polymerase (Quantabio), 11 μL of DNA template, 2.5 μL of 100 nM fluorescin, 1 μL of 25X primer, and TaqMan probe mix containing 25 μM primer and 6.25 μM of TaqMan probes (Supplementary Table 3). The reaction mixture volume was adjusted to a final volume of 27μL with Milli-Q. Samples were loaded into Sapphire chip slots (Stilla) and chips were placed into the Naica Geode (Stilla). The Naica Geode system was pressurized to 1.3 bar and samples were partitioned in approximately 25.000 droplets per chip. For amplification we used the following two-step PCR protocol: 10 minutes at 95°C, 45 cycles of 30 seconds at 95°C and 15 seconds at 58°C or 60°C (depending on primer/probe mixture used). Sapphire chips were removed from the Naica Geode and transferred to the Naica Prism3 (Stilla) for digital counting of the crystal droplets with inversions.

### cdPCR inversion detection

Chips were loaded into the Naica Prism3, using the Crystal Reader software (Stilla). The preset FAM and Cy-5 excitation wavelengths were selected. Exposure times for FAM and Cy-5 were set to 65ms and 50ms, respectively. Crystal droplet matrixes were inspected using the “Quality Control” section in the Crystal Reader software where image sharpness, number of saturated objects, and droplet number were assessed. Samples with fewer than 19.500 droplets were excluded from analysis. In the “Plots and Populations” section, crystal droplets were clustered and visualized as dots based on fluorescence intensity. By using the “2D dot plot” function, the lower-bound thresholds were set so that for both the FAM and the Cy-5 channel, the background fluorescence signals could be excluded. Per Cy-5 – FAM probe combination, it was confirmed that no bleed-through of Cy-5 signal was present in the FAM channel, and vice versa. Dots above these thresholds were counted as initial positive events. It was manually verified that these individual positive crystal droplets were above the background signal threshold in both the FAM and the Cy-5 channel by utilizing the ”Quality Control” section. In case a droplet showed only the FAM signal or only the Cy-5 signal, the droplet was regarded as not containing an inversion. However, in 99.9 % of the droplets both signals were present, or both were absent.

### Genome copy number estimations

For the quantification of the frequencies of inversions in the cdPCR system, we estimated the genome copy number per crystal droplet by multiplying the total amount of DNA in the sample (ng) with Avogadro’s constant and dividing the outcome with the genome size of tomato (bp) multiplied with the average base pair weight (Da) and with a 1 * 10^9^ gram- to-ng conversion.

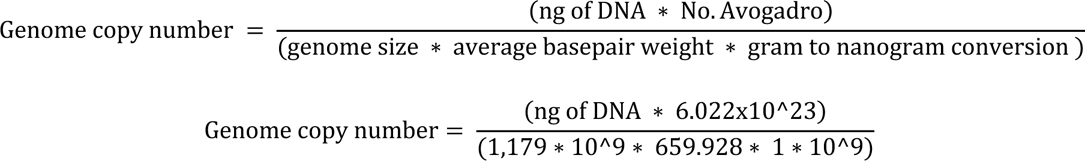

### CRISPR/Cas9 mutation induction efficiency analysis

In view of evaluation of the cutting efficiencies of the used CRISPR/Cas constructs, regions around the CRISPR/Cas9 target sites were amplified via PCR using Phusion polymerase (ThermoFisher) and primers containing an eight-base-pair barcode (Supplementary figure 3). The resulting amplicons were cleaned using the DNA Clean & Concentrator -5 kit (Zymo Research). Samples were pooled equimolarly and Illumina Hi-Seq sequenced (Eurofins Europe) followed by demultiplexing and trimming in Geneious Prime (https://www.geneious.com/prime/), for analysis of the mutations at the cutting sites.

CRISPR/Cas9 editing frequencies were acquired from the Hi-Seq data using the “amplicanPipeline” function in AmpliCan (Labun *et al.,* 2019; https://github.com/valenlab/amplican; Supplementary code 2). Analysis was performed using both reads per read pair. In the Amplican configuration file, a search window is required in which the script will search for mutations. In the configuration file we indicated the two nucleotides with the predicted CRISPR/Cas9 cutting site, using a 4 bp cut-off. Per analysed CRISPR/Cas9 site, two negative control samples were used (Supplementary code 1).

### PacBio Sequel II read inversion analysis

DNA fragments including the inversion events were amplified via PCR by utilizing site specific primers (Supplementary Table 3). For the 1 kb inversion samples, a 1758 bp amplicon was amplified, and for the 3 kb inversion an amplicon was generated of 3929 bp in size (Supplementary Table 4). Three replications were used per inversion length. DNA samples were prepared for PacBio SMRT-Bell sequencing by following the standard PacBio sequencing protocol (https://www.pacb.com/wp-content/uploads/Procedure-Checklist-Preparing-SMRTbell-Libraries-using-PacBio-Barcoded-Overhang-Adapters-for-Multiplexing-Amplicons.pdf). PacBio sequencing was facilitated by the Leids University Medical Center, The Netherlands.

Inversion events were identified in PacBio sequencing reads by using the “grep” command in Linux. At each DSB site in the amplicon, we searched for inversion events by specifying a 16 bp sequence at the left site and at the right site of the DSB location. These 16 bp sections were located approximately 60 bp away from each other, while the DSB site was directly in between the two 16 bp sections, allowing for small deletions at the DSB sites to be detected. In the grep command, we allowed for a random insertion of DNA sequence of 1 to 80 bp in length, to enable the detection of inversion events which were coupled with small insertions at the DSB sites (Supplementary code 2).

## Acknowledgements

We would like to express our gratitude to the following individuals and organizations for their contributions to this research. First, we thank Marga van Gent-Pelzer from the Biointeractions and Plant Health group at Wageningen University & Research for her expert assistance with the crystal digital PCR system. Our appreciation also goes to Stilla Technologies for providing the crystal digital PCR system used in this study.

We are grateful to Rolf Vossen and Susan Kloet from the Leiden Genome Technology Center for their support in generating the Hi-Seq and PacBio Sequel II sequencing data. Additionally, we would like to acknowledge Jan Schaart for his guidance in the establishment of the tomato protoplast transfection protocol.

Finally, we extend our appreciation to the Nederlandse Organisatie voor Wetenschappelijk Onderzoek (NWO) and Dutch Topsector Horticulture & Starting Materials for funding this research project. We also acknowledge the essential support from our industry partners: BASF Nunhems, Bejo Zaden, HZPC, KWS, SESVanderHave, and Syngenta.

- **Data Availability Statement:** Data supporting the findings of this study will be openly available in accordance with FAIR (Findable, Accessible, Interoperable, and Reusable) principles.
- **Funding Statement:** This research was supported by funding companies BASF Nunhems, Bejo Zaden, HZPC, KWS, SESVanderHave, and Syngenta, and also received funding from the NWO Graduate School Groene Topsectoren call, and the Dutch Topsector Horticulture & Starting Materials.
- **Conflict of Interest Disclosure:** The authors declare no conflict of interest regarding the publication of this article. This research was supported by BASF Nunhems, Bejo Zaden, HZPC, KWS, SESVanderHave, Syngenta, the NWO Graduate School Groene Topsectoren, and the Dutch Topsector Horticulture & Starting Materials. These funding sources had no role in the design, execution, interpretation, or writing of the study.
- Permission to Reproduce Material from Other Sources: Not applicable.

## Notes

### Competing Interest Statement

The authors have declared no competing interest.

