## Supplementary Information for "Key determinants of CRISPR/Cas9 induced inversions in tomato"

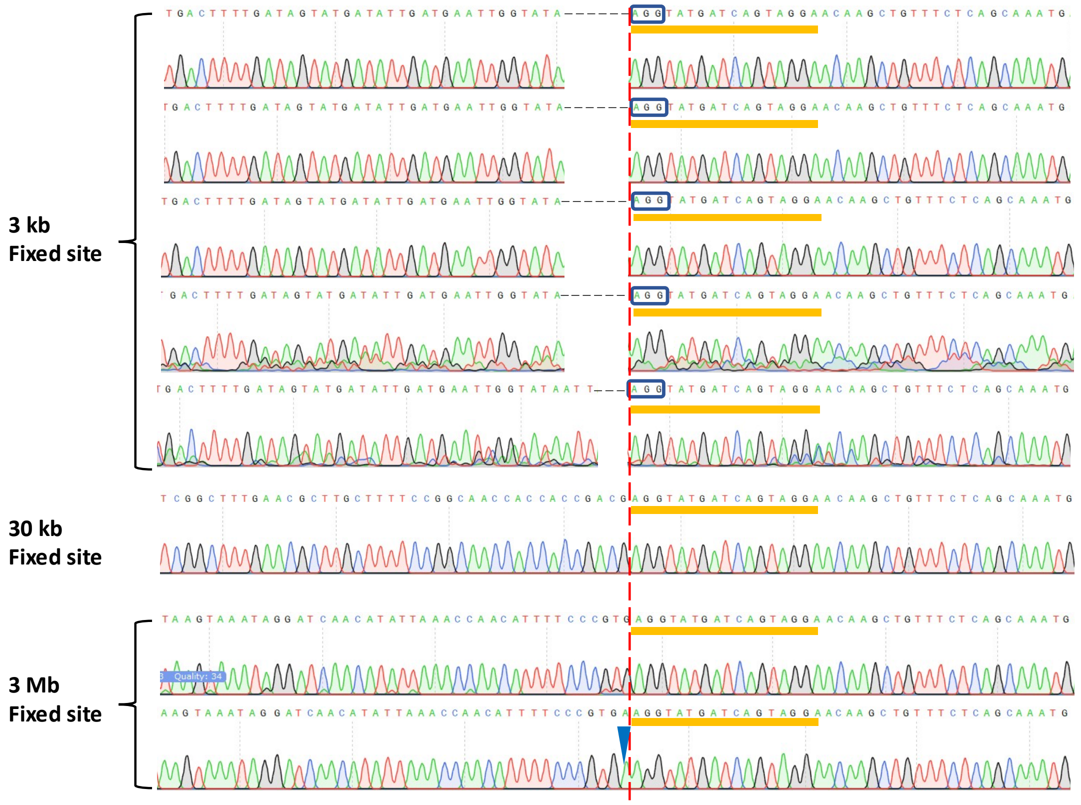

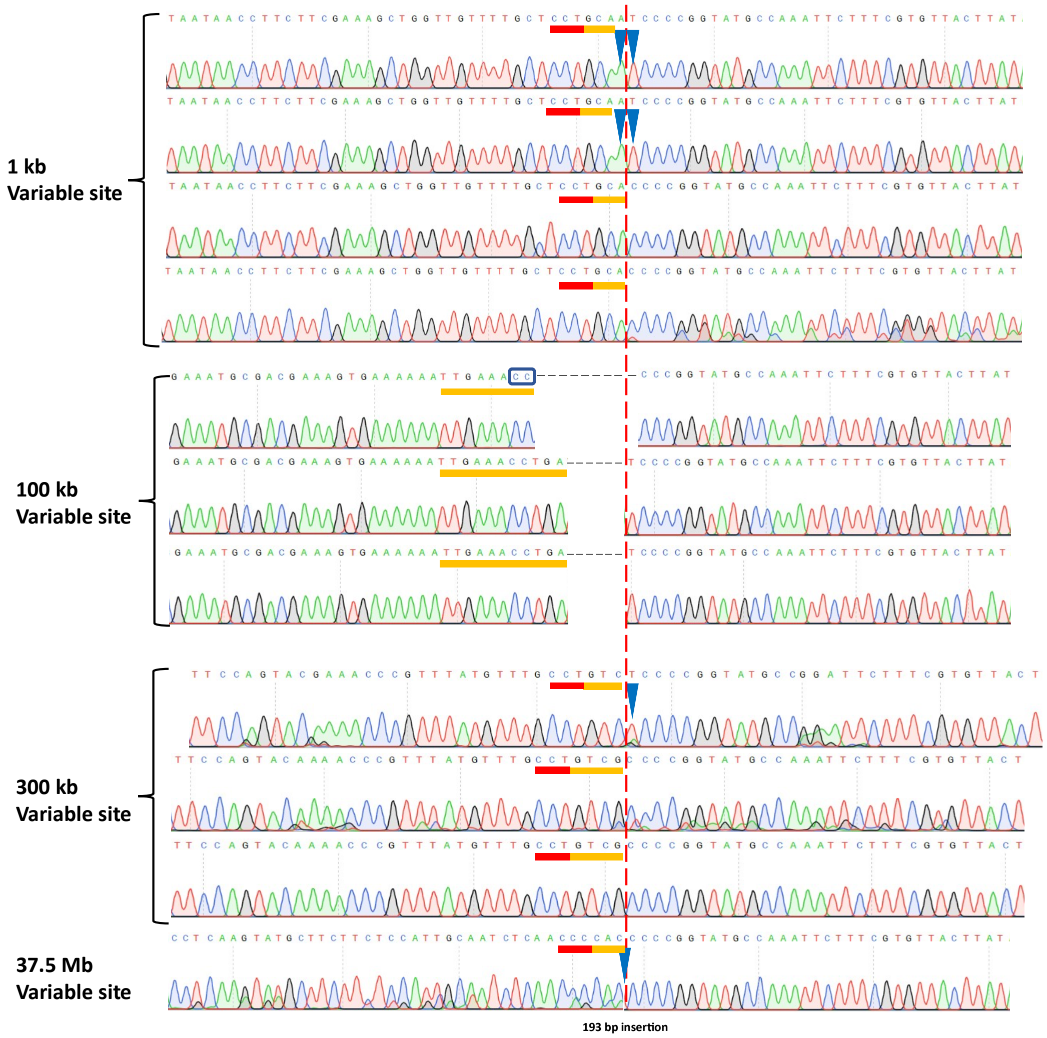


Supplementary figure 1. Cartoon depicting the sequences inversion borders at either the fixed- or the variable DSB site. Dashed Red line indicates the predicted DSB border. Black vertical dashes and black triangles indicate single base-pair deletions or insertions, respectively. Blue rectangle borders indicate suspected MMEJ-based repair patterns. gRNA sequences and corresponding PAM sites are depicted as orange and red bars above the Sanger sequences, respectively.

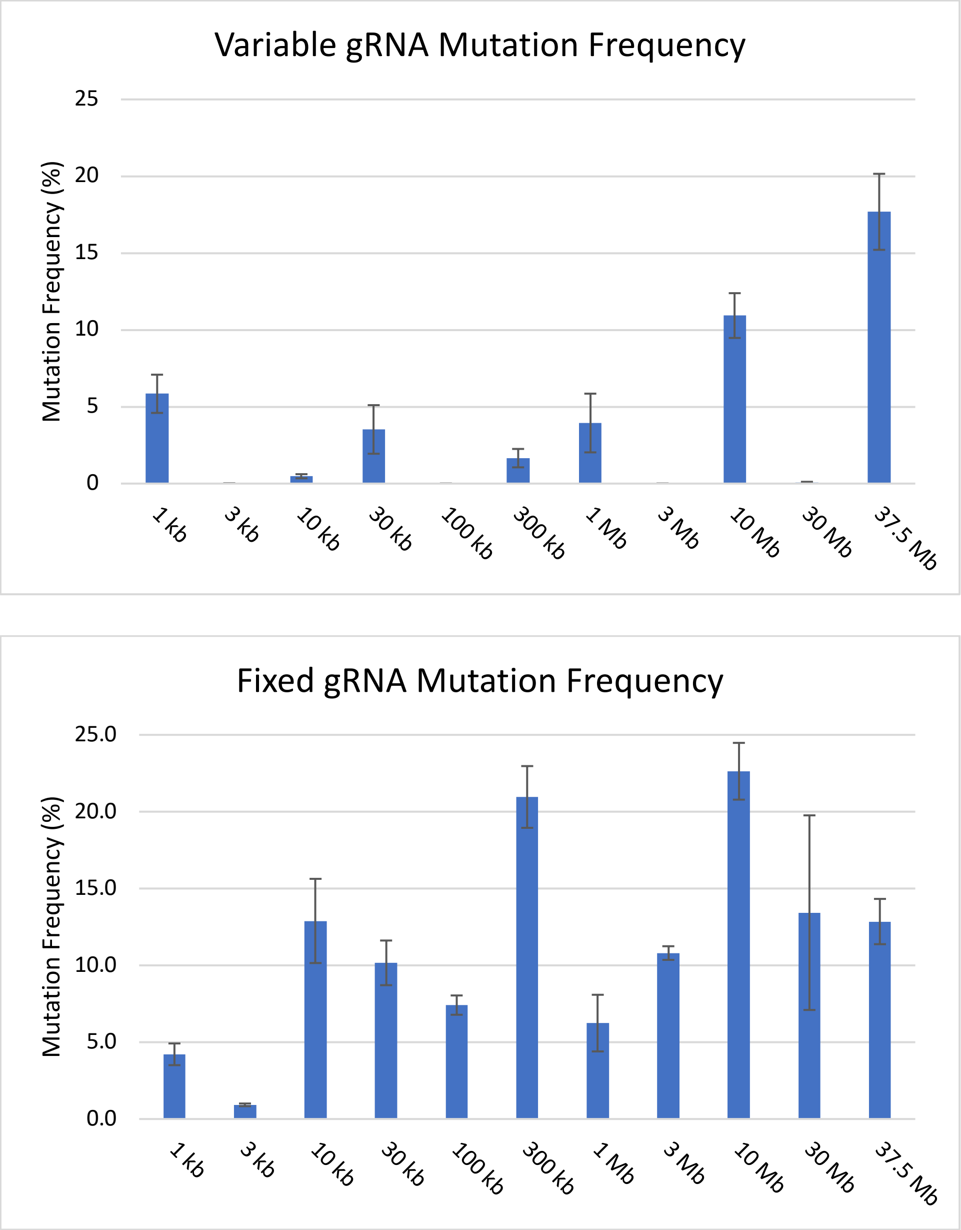


Supplementary Figure 2. Mutation frequencies in samples of 1 kb to 37.5 Mb in size for the variable gRNA (top) and for the fixed gRNA (bottom). Error bars show standard deviation.

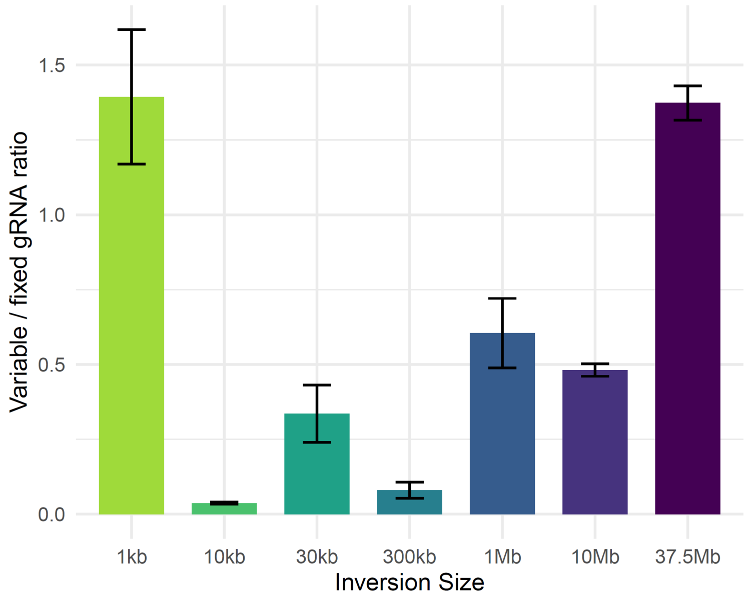


Supplementary figure 3. The ratio between the 'variable' and the 'fixed' gRNA mutation induction frequency for each inversion size. Error bars depict standard deviation.

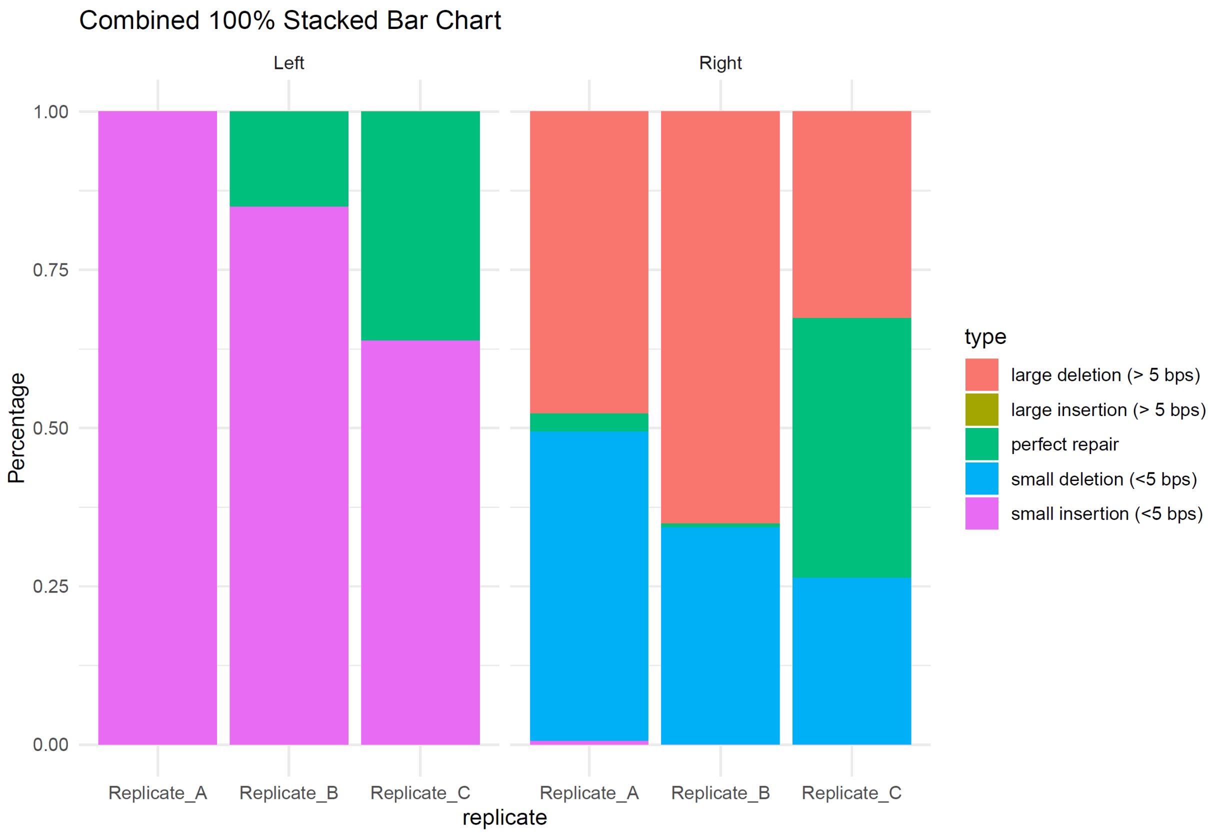


Supplementary figure 4. Analysis of sequence types at left and right double-strand break (DSB) sites across three replicates. The 100% stacked bar charts, display the distribution of different sequence types found at both the left and right DSB sites, being large deletions (> 5 bp), large insertions (> 5 bp), perfect repair, small deletions (< 5 bp), and small insertions (< 5 bp). The y-axis represents the percentage of reads for each sequence type. The bar charts are organized into two sets of three replicates each, labelled as A, B, and C. The first set of bar charts on the left shows the distribution of sequence types at the left DSB site, while the second set on the right displays the sequence types at the right DSB site.

### Supplementary Table 1

*Inversion induction gRNAs*

| gRNA | strand | gRNA name | Distance from "Fixed gRNA" (bp) | Inversions detected via Sanger sequencing? | Expected DSB position in SL4.0ch06 |
|---|---|---|---|---|---|
| TCCTACTGATCATACCTCCC**CGG** | - | Fixed gRNA | 0 | | 40808481 – 2 |
| TGTACAGGACAGAGCATTGC**AGG** | - | 1 kb | 1005 | Yes | 40807476 – 7 |
| AATTGGTATAATTAGGGAT**AGG** | - | 3 kb | 3135 | Yes | 40805346 – 7 |
| CAGGGAGTTGGTGTCTCCCC**TGG** | + | 10 kb | 10184 | No | 40798297 – 8 |
| TACTCCGTAACGTCCCCGT**CGG** | + | 30 kb | 29952 | Yes | 40778529 – 0 |
| TTGAAACCTGAGCAGCTGGG**AGG** | + | 100 kb | 99954 | Yes | 40708527 – 8 |
| AGTGAGTTACAGATTACGAC**AGG** | - | 300 kb | 297704 | Yes | 40510777 – 8 |
| TGCAATCTGCACAGGACCA**TGG** | - | 1 Mb | 994264 | No | 39814217 – 8 |
| TCATGGGTGGCATATGTCAC**GGG** | + | 3 Mb | 2986912 | Yes | 37821569 – 0 |
| AGATCTGTGTCGTTGCTCCA**TGG** | + | 10 Mb | 9873024 | No | 30935457 – 8 |
| AGGAACGGACTGGGCTCAAG**CGG** | - | 30 Mb | 29395350 | No | 11413131 – 2 |
| TCGTCCATCTTATGGCAGTG**GGG** | - | 37.5 Mb | 37509359 | Yes | 3299122 – 3 |

**Supplementary Table 2**

| Vector component | Sequence |
|---|---|
| NosP::NPTII | Gaaccgcaacgttgaaggagccactgagccgcgggtttctggagtttaatgagctaagcacatacgtcagaaaccattattgcgcgt<br>Tcaaaagtcgcctaaggtcactatcagctagcaaatatttcttgtcaaaaatgctccactgacgttccataaattcccctcggtatcca<br>Attagagtctcatattcactctcctatttttacaacaattaccaacaacaacaaacaacaaacaacattacaattacatttacaattac<br>Catggttgaacaagatggattgcacgcaggttctccggccgcttgggtggagaggctattcggctatgactgggcacaacagacaat<br>Cggctgctctgatgccgccgtgttccggctgtcagcgcaggggcgcccggttctttttgtcaagaccgacctgtccggtgccctgaatg<br>Aactgcaggacgaggcagcgcggctatcgtggctggccacgacgggcgttccttgcgcagctgtgctcgacgttgtcactgaagcgg<br>Gaagggactggctgctattgggcgaagtgccggggcaggatctcctgtcatctcaccttgctcctgccgagaaagtatccatcatggc<br>Tgatgcaatgcggcggctgcatacgcttgatccggctacctgcccattcgaccaccaagcgaaacatcgcatcgagcgagcacgtac<br>Tcggatggaagccggtcttgtcgatcaggatgatctggacgaagagcatcaggggctcgcgccagccgaactgttcgccaggctcaa<br>Ggcgcgcatgcccgacggcgaggatctcgtcgtgactcatggcgatgcctgcttgccgaatatcatggtggaaaatggccgcttttctg<br>Gattcatcgactgtggccggctgggtgtggcggaccgctatcaggacatagcgttggctacccgtgatattgctgaagagcttggcggc<br>Gaatgggctgaccgcttcctcgtgctttacggtatcgccgctcccgattcgcagcgcatcgccttctatcgccttcttgacgagttcttctg<br>A |
| pUbi::Cas9 | |
| pU6:26::gRNA | |
| pCsVMV::GFP | |

**Supplementary Table 3**

| Name | Sequence |
|---|---|
| cdPCR_Fixed_RV | GCATGGAGAGGATACTTGAAAGA |
| cdPCR_1kb_RV | TTCCAACAAGTCTTGCAATCA |
| cdPCR_3kb_RV | ACGAATGCCAGCAAGAATTT |
| cdPCR_10kb_RV | TTAGCAGATATCAGGGGCAAT |
| cdPCR_30kb_RV | ACTTAGGTTCCTCCGGTGTG |
| cdPCR_100kb_RV | AAGAGTTGTATGGCAACTTTCAGA |
| cdPCR_Hiseq_300kb_RV_v2 | GAACTAGAAGCATCGTATGAATGG |
| cdPCR_1000kb_RV | AAGGTTCTTTACCCGTCTGATG |
| cdPCR_3000kb_RV | TGGAATTGTGACGTGATATGC |
| cdPCR_10000kb_RV | TGGATGGCAATACATTAGGACAA |
| cdPCR_30000kb_RV | TGGTACAACAAATGCAAGTAACTG |
| cdPCR_Hiseq_37500kb_RV_v2 | GGATGTTCAAATCACTCTATGTGG |
| "Fixed" Probe | TCAGTAGGAACAAGCTGTTTCTCAGCA |
| 1 kb Probe | CCTGAGTCTAACACAACCTCAAAAGC |
| 10 kb Probe | TGTGAGACTAGTCACCGCTCTA |
| 30 kb Probe | CGGAAAAGCAAGCGTTCAAAGC |
| 100 kb Probe | CAAATGATTCCTCGAGGCAAACCTGC |
| 300 kb Probe | CACTTTGGTACCTTACCACTGAAACC |
| 1000 kb Probe | ATGTTTTTCTTTTCAGTGCAATCTGCAC |
| 3000 kb Probe | ATCAACATATTAAACCAACATTTTCCCG |
| 10000 kb Probe | CATCGATCTGACAATTGATTCATCAAG |
| 30000 kb Probe | CAGTCCGTTCCTTTCTCAGATGAATAAAC |
| 37500 kb Probe | CCTTTGTTTTGCTAACGTATCGTATC |
| Fixed_FW_1kb | AACCAAGGAACTTCCTCCTCAAAAACGAGA |
| Fixed_FW_3kb | AAGGTACGAACTTCCTCCTCAAAAACGAGA |
| Fixed_FW_10kb | ACCTACCTAACTTCCTCCTCAAAAACGAGA |
| Fixed_FW_30kb | ACGTGTTGAACTTCCTCCTCAAAAACGAGA |
| Fixed_FW_100kb | ACTGGACTAACTTCCTCCTCAAAAACGAGA |
| Fixed_FW_300kb | AGAGACTGAACTTCCTCCTCAAAAACGAGA |
| Fixed_FW_1000kb | AGTCGACTAACTTCCTCCTCAAAAACGAGA |
| Fixed_FW_3000kb | ATATGCCGAACTTCCTCCTCAAAAACGAGA |
| Fixed_FW_10000kb | CAACCATGAACTTCCTCCTCAAAAACGAGA |
| Fixed_FW_30000kb | CACAGTGTAACTTCCTCCTCAAAAACGAGA |
| Fixed_FW_37500kb | CAGAAGTGAACTTCCTCCTCAAAAACGAGA |
| Fixed_FW_Control | CAGTGACTAACTTCCTCCTCAAAAACGAGA |
| Fixed_RV_Rep1 | AACCGGTTTATGTATGTAATTAGCATGGAGAGG |
| Fixed_RV_Rep2 | ACACACTGTATGTATGTAATTAGCATGGAGAGG |
| Fixed_RV_Rep3 | ACCTAGGTTATGTATGTAATTAGCATGGAGAGG |
| Fixed_RV_Rep4 | ACGTTGGTTATGTATGTAATTAGCATGGAGAGG |
| 1kb_FW_Treatm | ACTGTCAGCAAGTAGATGTATTTTATTTGGGGATA |
| 1kb_FW_Control | AGGAGAAGCAAGTAGATGTATTTTATTTGGGGATA |
| 3kb_FW_Treatm | ACTGTCAGACCTTTTCGAATAGTTTAGGGATATT |
| 3kb_FW_Control | AGGAGAAGACCTTTTCGAATAGTTTAGGGATATT |
| 10kb_FW_Treatm | ACTGTCAGCAGTTAACCTTCATGTTTCACTTCC |
| 10kb_FW_Control | AGGAGAAGCAGTTAACCTTCATGTTTCACTTCC |

| | |
|---|---|
| 30kb_FW_Treatm | ACTGTCAGTCTCAGCCACGTCACTTCTG |
| 30kb_FW_Control | AGGAGAAGTCTCAGCCACGTCACTTCTG |
| 100kb_FW_Treatm | ACTGTCAGTTGGACCGTGATCTAAAAAGTTC |
| 100kb_FW_Control | AGGAGAAGTTGGACCGTGATCTAAAAAGTTC |
| 300kb_FW_Treatm | ACTGTCAGTTGGTCTTACAAATGAAGAGAAAAAT |
| 300kb_FW_Control | AGGAGAAGTTGGTCTTACAAATGAAGAGAAAAAT |
| 1000kb_FW_Treatm | ACTGTCAGCAGAACAATTGAAGTTGGATATCCC |
| 1000kb_FW_Control | AGGAGAAGCAGAACAATTGAAGTTGGATATCCC |
| 3000kb_FW_Treatm | ACTGTCAGCGAACTTCTCTATATTCACCGCTAT |
| 3000kb_FW_Control | AGGAGAAGCGAACTTCTCTATATTCACCGCTAT |
| 10000kb_FW_Treatm | ACTGTCAGAACTTCTCTGCTTATTGGGTTCA |
| 10000kb_FW_Control | AGGAGAAGAACTTCTCTGCTTATTGGGTTCA |
| 30000kb_FW_Treatm | ACTGTCAGTTCCATGAGACAAGCTTTATGGT |
| 30000kb_FW_Control | AGGAGAAGTTCCATGAGACAAGCTTTATGGT |
| 37500kb_FW_Treatm | ACTGTCAGTCTTCATTATAGTCTTCCTCACCTCA |
| 37500kb_FW_Control | AGGAGAAGTCTTCATTATAGTCTTCCTCACCTCA |
| 1kb_RV_Rep1 | AACCGGTTGAGGTTGTGTTAGACTCAGGTTCA |
| 1kb_RV_Rep2 | ACACACTGGAGGTTGTGTTAGACTCAGGTTCA |
| 1kb_RV_Rep3 | ACCTAGGTGAGGTTGTGTTAGACTCAGGTTCA |
| 1kb_RV_Rep4 | ACGTTGGTGAGGTTGTGTTAGACTCAGGTTCA |
| 3kb_RV_Rep1 | AACCGGTTACGAATGCCAGCAAGAATTT |
| 3kb_RV_Rep2 | ACACACTGACGAATGCCAGCAAGAATTT |
| 3kb_RV_Rep3 | ACCTAGGTACGAATGCCAGCAAGAATTT |
| 3kb_RV_Rep4 | ACGTTGGTACGAATGCCAGCAAGAATTT |
| 10kb_RV_Rep1 | AACCGGTTGGGCAATAGTAGAGCGGTGA |
| 10kb_RV_Rep2 | ACACACTGGGGCAATAGTAGAGCGGTGA |
| 10kb_RV_Rep3 | ACCTAGGTGGGCAATAGTAGAGCGGTGA |
| 10kb_RV_Rep4 | ACGTTGGTGGGCAATAGTAGAGCGGTGA |
| 30kb_RV_Rep1 | AACCGGTTACTTAGGTTCCTCCGGTGTG |
| 30kb_RV_Rep2 | ACACACTGACTTAGGTTCCTCCGGTGTG |
| 30kb_RV_Rep3 | ACCTAGGTACTTAGGTTCCTCCGGTGTG |
| 30kb_RV_Rep4 | ACGTTGGTACTTAGGTTCCTCCGGTGTG |
| 100kb_RV_Rep1 | AACCGGTTCAACTTTCAGATATTGTCAAATGATTCC |
| 100kb_RV_Rep2 | ACACACTGCAACTTTCAGATATTGTCAAATGATTCC |
| 100kb_RV_Rep3 | ACCTAGGTCAACTTTCAGATATTGTCAAATGATTCC |
| 100kb_RV_Rep4 | ACGTTGGTCAACTTTCAGATATTGTCAAATGATTCC |
| 300kb_RV_Rep1 | AACCGGTTCATCGTATGAATGGCAAAGC |
| 300kb_RV_Rep2 | ACACACTGCATCGTATGAATGGCAAAGC |
| 300kb_RV_Rep3 | ACCTAGGTCATCGTATGAATGGCAAAGC |
| 300kb_RV_Rep4 | ACGTTGGTCATCGTATGAATGGCAAAGC |
| 1000kb_RV_Rep1 | AACCGGTTGTTTCAAACAGATTTCATATTTCATTT |
| 1000kb_RV_Rep2 | ACACACTGGTTTCAAACAGATTTCATATTTCATTT |
| 1000kb_RV_Rep3 | ACCTAGGTGTTTCAAACAGATTTCATATTTCATTT |
| 1000kb_RV_Rep4 | ACGTTGGTGTTTCAAACAGATTTCATATTTCATTT |
| 3000kb_RV_Rep1 | AACCGGTTTGGAATTGTGACGTGATATGC |
| 3000kb_RV_Rep2 | ACACACTGTGGAATTGTGACGTGATATGC |
| 3000kb_RV_Rep3 | ACCTAGGTTGGAATTGTGACGTGATATGC |
| 3000kb_RV_Rep4 | ACGTTGGTTGGAATTGTGACGTGATATGC |

| | |
|---|---|
| 10000kb_RV_Rep1 | AACCGGTTTGAGAATGATGGACAATTAAGAACA |
| 10000kb_RV_Rep2 | ACACACTGTGAGAATGATGGACAATTAAGAACA |
| 10000kb_RV_Rep3 | ACCTAGGTTGAGAATGATGGACAATTAAGAACA |
| 10000kb_RV_Rep4 | ACGTTGGTTGAGAATGATGGACAATTAAGAACA |
| 30000kb_RV_Rep1 | AACCGGTTTGGTACAACAAATGCAAGTAACTG |
| 30000kb_RV_Rep2 | ACACACTGTGGTACAACAAATGCAAGTAACTG |
| 30000kb_RV_Rep3 | ACCTAGGTTGGTACAACAAATGCAAGTAACTG |
| 30000kb_RV_Rep4 | ACGTTGGTTGGTACAACAAATGCAAGTAACTG |
| 37500kb_RV_Rep1 | AACCGGTTCATGCCCCTCACAAATGC |
| 37500kb_RV_Rep2 | ACACACTGCATGCCCCTCACAAATGC |
| 37500kb_RV_Rep3 | ACCTAGGTCATGCCCCTCACAAATGC |
| 37500kb_RV_Rep4 | ACGTTGGTCATGCCCCTCACAAATGC |

### Supplementary Table 4

*PacBio sequencing amplicon sizes*

| Fragment | Range SL4.0ch06 | Real size |
|---|---|---|
| **1 kb WT** | 40807126..40808883 | 1,758 bp |
| **3 kb WT** | 40804955..40808883 | 3,929 bp |

### Supplementary code 1

```
# Install amplican (Make sure you have the latest version of R first)

if (!requireNamespace("BiocManager", quietly = TRUE))
  install.packages("BiocManager")

BiocManager::install("amplican")

# Load amplican:

library(amplican)

## Run amplicanPileline:
amplicanPipeline(config,
        fastq_folder,
        results_folder,
        knit_reports = TRUE,
        write_alignments_format = "txt",
        average_quality = 30, min_quality = 5,
        use_parallel = TRUE,
        scoring_matrix = Biostrings::nucleotideSubstitutionMatrix(match = 5,
                                        mismatch = -4,
                                        baseOnly = TRUE,
                                        type = "DNA"),
        gap_opening = 25,
        gap_extension = 0,
        fastqfiles = 0,
        primer_mismatch = 0,
        donor_mismatch = 3,
        PRIMER_DIMER = 30,
        event_filter = TRUE,
        cut_buffer = 4,
        promiscuous_consensus = TRUE,
        normalize = c("guideRNA", "Group"))
```

### Supplementary code 2

### Read search left site of the DSB, FW and RV

Reads

```
grep -c "GAACCTCAAGTCTGGC.\{1,80\}TATATATATATCGTCA\|TGACGATATATATATA.\{1,80\}GCCAGACTTGAGGTTC" 100bp-INV-a.fasta
grep -c "AGAGCTGAATCAAGCT.\{1,80\}TATATATATATCGTCA\|TGACGATATATATATA.\{1,80\}AGCTTGATTCAGCTCT" 300bp-INV-a.fasta
grep -c "GATATTGTAATAACCT.\{1,80\}TATATATATATCGTCA\|TGACGATATATATATA.\{1,80\}AGGTTATTACAATATC" 1000bp-INV-a.fasta
grep -c "TGAACCTTTTCGAATA.\{1,80\}TATATATATATCGTCA\|TGACGATATATATATA.\{1,80\}TATTCGAAAAGGTTCA" 3000bp-INV-a.fasta
```

### Read search right site of the DSB, FW and RV

Reads

```
grep -c "AATCTCGTTTTTGAGG.\{1,80\}CAGCAAATGAGTATCT\|AGATACTCATTTGCTG.\{1,80\}CCTCAAAAACGAGATT" 100bp-INV-a.fasta
grep -c "TGATATGTTTGCAGGC.\{1,80\}CAGCAAATGAGTATCT\|AGATACTCATTTGCTG.\{1,80\}GCCTGCAAACATATCA" 300bp-INV-a.fasta
grep -c "AAATTGAGTAACTTTT.\{1,80\}CAGCAAATGAGTATCT\|AGATACTCATTTGCTG.\{1,80\}AAAAGTTACTCAATTT" 1000bp-INV-a.fasta
grep -c "TATTTGACTTTTGATA.\{1,80\}CAGCAAATGAGTATCT\|AGATACTCATTTGCTG.\{1,80\}TATCAAAAGTCAAATA" 3000bp-INV-a.fasta
```

### Read search left and right site of the DSB, FW and RV

Reads

```
grep -B 1
"GAACCTCAAGTCTGGC.\{1,80\}TATATATATATCGTCA\|TGACGATATATATATA.\{1,80\}GCCAGACTTGAGGTTC\|AATCTCGTTTTTGAGG.\{1,80\}
CAGCAAATGAGTATCT\|AGATACTCATTTGCTG.\{1,80\}CCTCAAAAACGAGATT" 100bp-INV-a.fasta > 100bp-INV-a-Tot.fasta
grep -B 1
"AGAGCTGAATCAAGCT.\{1,80\}TATATATATATCGTCA\|TGACGATATATATATA.\{1,80\}AGCTTGATTCAGCTCT\|TGATATGTTTGCAGGC.\{1,80\}
CAGCAAATGAGTATCT\|AGATACTCATTTGCTG.\{1,80\}GCCTGCAAACATATCA" 300bp-INV-a.fasta > 300bp-INV-a-Tot.fasta
grep -B 1
"GATATTGTAATAACCT.\{1,80\}TATATATATATCGTCA\|TGACGATATATATATA.\{1,80\}AGGTTATTACAATATC\|AAATTGAGTAACTTTT.\{1,80\}C
AGCAAATGAGTATCT\|AGATACTCATTTGCTG.\{1,80\}AAAAGTTACTCAATTT" 1000bp-INV-a.fasta > 1000bp-INV-a-Tot.fasta
grep -B 1
"TGAACCTTTTCGAATA.\{1,80\}TATATATATATCGTCA\|TGACGATATATATATA.\{1,80\}TATTCGAAAAGGTTCA\|TATTTGACTTTTGATA.\{1,80\}C
AGCAAATGAGTATCT\|AGATACTCATTTGCTG.\{1,80\}TATCAAAAGTCAAATA" 3000bp-INV-a.fasta > 3000bp-INV-a-Tot.fasta
```

### Read search right site of the DSB, FW and RV, using only the files with a “left from DSB site” match

```
grep -B 1 "GAACCTCAAGTCTGGC.\{1,80\}TATATATATATCGTCA\|TGACGATATATATATA.\{1,80\}GCCAGACTTGAGGTTC" 100bp-INV-a-
Tot.fasta > 100bp-INV-a-L.fasta
grep -B 1 "AGAGCTGAATCAAGCT.\{1,80\}TATATATATATCGTCA\|TGACGATATATATATA.\{1,80\}AGCTTGATTCAGCTCT" 300bp-INV-a-
Tot.fasta > 300bp-INV-a-L.fasta
grep -B 1 "GATATTGTAATAACCT.\{1,80\}TATATATATATCGTCA\|TGACGATATATATATA.\{1,80\}AGGTTATTACAATATC" 1000bp-INV-a-
Tot.fasta > 1000bp-INV-a-L.fasta
grep -B 1 "TGAACCTTTTCGAATA.\{1,80\}TATATATATATCGTCA\|TGACGATATATATATA.\{1,80\}TATTCGAAAAGGTTCA" 3000bp-INV-a-
Tot.fasta > 3000bp-INV-a-L.fasta
```
